# Dexmedetomidine - commonly used in functional imaging studies - increases susceptibility to seizures in rats but not in wild type mice

**DOI:** 10.1101/2019.12.29.890525

**Authors:** Aleksandra Bortel, Roland Pilgram, Ze Shan Yao, Amir Shmuel

## Abstract

Functional MRI (fMRI) utilizes changes in metabolic and hemodynamic signals to indirectly infer the underlying local changes in neuronal activity. To investigate the mechanisms of fMRI responses, spontaneous fluctuations, and functional connectivity in the resting-state, it is important to pursue fMRI in animal models. Animal studies commonly use dexmedetomidine sedation. It has been demonstrated that potent sensory stimuli administered under dexmedetomidine are prone to inducing seizures in Sprague-Dawley (SD) rats.

Here we combined optical imaging of intrinsic signals and cerebral blood flow with neurophysiological recordings to measure responses in rat area S1FL to electrical forepaw stimulation administered at 8 Hz. We show that the increased susceptibility to seizures starts no later than 1 hour and ends no sooner than 3 hours after initiating a continuous administration of dexmedetomidine. By administering different combinations of anesthetic and sedative agents, we demonstrate that dexmedetomidine is the sole agent necessary for the increased susceptibility to seizures. The increased susceptibility to seizures prevails under a combination of 0.3%-0.5% isoflurane and dexmedetomidine anesthesia. The blood-oxygenation and cerebral blood flow responses to seizures induced by forepaw stimulation have a higher amplitude and a larger spatial extent relative to physiological responses to the same stimuli. The epileptic activity and the associated blood oxygenation and cerebral blood flow responses stretched beyond the stimulation period. We observed seizures in response to forepaw stimulation with 1-2 mA pulses administered at 8 Hz. In contrast, responses to stimuli administered at 4 Hz were seizure-free. We demonstrate that such seizures are generated not only in SD rats but also in Long-Evans rats, but not in C57BL6 mice stimulated with similar potent stimuli under dexmedetomidine sedation.

We conclude that high-amplitude hemodynamic functional imaging responses evoked by peripheral stimulation in rats sedated with dexmedetomidine are possibly due to the induction of epileptic activity. Therefore, caution should be practiced in experiments that combine the administration of potent stimuli with dexmedetomidine sedation. We propose stimulation paradigms that elicit seizure-free, well detectable neurophysiological and hemodynamic responses in rats. We further conclude that the increased susceptibility to seizures under dexmedetomidine sedation is species dependent.

## INTRODUCTION

Functional connectivity (FC) refers to the temporal correlation between spatially remote neurophysiological events (Friston et al., 1993). FC analysis based on functional magnetic resonance imaging (fMRI) makes it possible to obtain an approximation of the pattern of thalamocortical and cortico-cortical connections noninvasively; thus, it is readily usable on human subjects. FC analysis can be pursued using data obtained during subject stimulation, task performance, or in the resting-state. In addition to shedding light on the pattern of connections, FC carries information that can be used to detecting a malfunction of the brain in disease (Fox and Greicius, 2010).

fMRI of the resting-state utilizes spontaneous fluctuations in metabolic and hemodynamic signals to infer the underlying local changes in neuronal activity (Shmuel and Leopold, 2008). Thus, the fMRI signal is an indirect measure of changes in neuronal activity. Therefore, for correct interpretation of spontaneous fluctuations and FC in the resting-state, it is important to characterize the neuronal mechanisms of these phenomena by combining fMRI and neurophysiology in animal models.

Animal studies of the resting-state and neurovascular coupling in general commonly use dexmedetomidine. Dexmedetomidine has been the sedative of choice for long functional imaging and neurophysiology studies in rats (Fukuda et al., 2013; Nasrallah et al., 2012; Pawela et al., 2008; Sotero et al., 2010; Zhao et al., 2008) and mice (Adamczak et al., 2010; Bukhari et al., 2017). It was previously reported that medetomidine sedation does not affect seizure vulnerability, nor does it affect LFP and BOLD responses during seizures evoked by systemic administration of kainic acid in rats (Airaksinen et al., 2012; Airaksinen et al., 2010). In contrast, Fukuda et al. (2013) demonstrated that sedation induced by intravenous administration of dexmedetomidine for more than two hours changes seizure susceptibility in rats. Forelimb stimulation elicited seizure-like responses accompanied by changes in cerebral blood flow (CBF) in Sprague-Dawley (SD) rats (Fukuda et al., 2013). These results have been corroborated by Bortel et al. (2019), who showed that seizures can be elicited not only with forelimb stimulation but also with the less-potent digit stimulation and that they propagated from the onset zone to adjacent cortical areas. Bortel et al. (2019) observed high-amplitude, high-frequency oscillations prior to and during the seizures, and demonstrated that the seizures are not induced by damage caused by inserting the electrodes into the cortex. To elicit the seizure, Fukuda et al. (2013) and Bortel et al. (2019) stimulated the forelimb or digit over 10 s long blocks. In contrast, short (e.g., 1 s) stimulation blocks of SD rats under dexmedtomidine sedation are not sufficient for inducing epileptiform activity (Sotero et al., 2015). Therefore, electrical stimuli administered at 8 Hz over a long duration under dexmedetomidine sedation elicit seizures.

Such potent forepaw stimulation - in the range of 9-12 Hz - are commonly used for imaging under medetomidine, isoflurane, or urethane anesthesia as these frequencies elicit the strongest BOLD response (Albers et al., 2018; Huttunen et al., 2008; Kim et al., 2010; Krautwald and Angenstein, 2012; Lambers et al., 2020; Masamoto et al., 2007; Nunes et al., 2019; Paasonen et al., 2017; Zhao et al., 2008). Therefore, it is important to test whether these stimuli induce seizures that could influence the hemodynamic responses. Moreover, mice sedated with dexmedetomidine are commonly used for studying the mechanisms underlying neurovascular coupling and resting-state fMRI. However, whether dexmedetomidine increases the susceptibility to seizures in mice remains unknown.

Here we demonstrate that the seizures induced by forepaws stimulation in SD rats are accompanied by spatially extended cerebral blood flow and blood oxygenation responses. We show that the increased susceptibility to seizures starts no later than 1 hour after initiating the dexmedetomidine administration and lasts for at least 2 hours. We demonstrate that the seizure susceptibility depends on the stimulation parameters and anesthesia regime. We propose reliable anesthesia regimes along with forepaw stimulation parameters that do not induce seizures but do generate well detectable hemodynamic responses. We further show that dexmedetomidine sedation increases the susceptibility to seizures not only in SD rats but also in Long-Evans (LE) rats. In contrast, C57BL6 (C57) wild type mice do not show susceptibility to seizures under the same anesthesia regime and stimulation protocol. We conclude that dexmedetomidine sedation increases the susceptibility to seizures accompanied by extended hemodynamic responses in SD and LE rats but not in wild-type mice.

## MATERIALS AND METHODS

### Animals, surgical procedures, and anesthesia

All procedures were approved by the animal care committees of the Montreal Neurological Institute and McGill University and were carried out in accordance with the guidelines of the Canadian Council on Animal Care. Here we report on experiments performed on 19 adult (100–107 days old) male SD rats weighing 440–560 g, 5 adult (100–109 days old) male LE rats weighing 475–495 g, and 5 age-matched adult C57 mice weighing 30-37g. In Table 1 – which summarizes the conditions under which we observed seizures – we include results obtained from 6 additional 100–107 days old male SD rats, on which we reported in detail in Bortel et al. (2019). All 25 SD rats, 5 LE rats, and 5 C57 mice were stimulated with electrical forepaw stimulation. The rats and mice were housed under the same controlled environmental conditions at 22±2° C with a 12h light/12h dark cycle (lights on from 7:00 a.m. to 7:00 p.m.) and received food and water ad libitum. A brief description of the surgical procedures is included below. All procedures carried out for experiments in SD rats are described in detail in Bortel et al. (2019). The procedures applied for experiments in LE rats were identical to those used for experiments in SD rats.

**Table 1.** Average number of rats and mice with seizures and average number of seizures per run according to anesthesia regime and stimulation parameters. The first column presents the species and strain (SD rats, LE rats or C57 mice) and the type of stimulation, i.e. forepaw stimulation. The second column presents the anesthesia regime, i.e. dexmedetomidine with buprenorphine sedation (**Dex & Bup**), dexmedetomidine with buprenorphine sedation without ketamine (**Dex & Bup without Ketamine**), and dexmedetomidine with isoflurane without buprenorphine (**Dex & Iso without Bup**). All neurophysiology data were recorded using a linear probe, except for the data presented in the second row labeled ‘**Dex & Bup measured with ECoG**‘ which were recorded using electrocorticography arrays. The experiments in this row, whose anesthesia regime is labeled in red font were presented in detail in Bortel et al. (2019). The 3^rd^, 5^th^, 7^th^, 9^th^, 11^th^ and 13^th^ columns present the number of rats/mice with seizures and the 4^th^, 6^th^, 8^th^, 10^th^, 12^th^ and 14^th^ columns present the average number of seizures per run (including all runs in rats and mice that had or did not have seizures). Results are expressed as a mean ± SEM.

| SPECIES and Stimulation TYPE | ANESTHESIA REGIME | STIMULATION PARAMETERS |  |  |  |  |  |  |  |  |  |  |  |
| --- | --- | --- | --- | --- | --- | --- | --- | --- | --- | --- | --- | --- | --- |
|  |  | 4Hz |  |  |  |  |  | 6Hz |  | 8Hz |  |  |  |
|  |  | 1 mA |  | 1.5 mA |  | 2 mA |  | 2 mA |  | 1 mA |  | 2 mA |  |
|  |  | Rats with seizures | No. of seizures per run | Rats with seizures | No. of seizures per run | Rats with seizures | No. of seizures per run | Rats with seizures | No. of seizures per run | Rats with seizures | No. of seizures per run | Rats with seizures | No. of seizures per run |
| **SD RATS**<br>Forepaw stim | **Dex & Bup** | 0/3 | – |  |  | 0/3 | – |  |  | 3/3 | 1.50 ±0.81 | 7/9 | 2.14 ±0.40 |
|  | **Dex & Bup measured with ECoG** |  |  |  |  |  |  |  |  |  |  | 5/6 | 3.04 ±0.42 |
|  | **Dex & Bup without Ketamine** |  |  |  |  |  |  |  |  |  |  | 3/3 | 2.37 ±0.35 |
|  | **Dex & Iso without Bup** |  |  | 0/3 | – |  |  |  |  |  |  | 2/4 | 1.88 ±0.44 |
| **LE RATS**<br>Forepaw stim | **Dex & Bup** |  |  |  |  |  |  |  |  |  |  | 5/5 | 3.73 ±0.50 |
| **C57 MICE**<br>Forepaw stim | **Dex & Bup** |  |  |  |  | 0/5 | – | 0/5 | – |  |  | 0/5 | – |

The rats were first injected with the anti-inflammatory drug carprofen (5 mg/kg SC; Zoetis, Canada) and anesthetized with a solution of xylazine (10 mg/kg IP; Bayer Inc., Canada) and ketamine (50 mg/kg IP; Wyeth, Canada). They were then intubated and placed in a stereotaxic frame. The surgical procedure was performed under ventilation with 100% oxygen and anesthesia with isoflurane (0.6-2%; Benson Medical Industries Inc., Canada). The scalp was incised to expose the skull covering the primary somatosensory cortex (S1) of the left hemisphere. One stainless steel screw (2.4 mm in length) – used as a ground and reference – was fixed to the skull above the visual cortex of the right hemisphere. The part of the skull overlying S1 was thinned until soft and transparent. We then performed an approximate 4 mm wide square-shaped craniotomy, centered on the forelimb representation in area S1FL based on a stereotaxic atlas (AP 0.00 mm, ML ±4.50 mm, DV −2.5 mm; Paxinos and Watson (2005)). The dura mater within the craniotomy was resected. At the end of the surgical procedure, a silicon chamber was created around the craniotomy and filled with Hank’s Balanced Salt Solution (Invitrogen, Canada).

Following the surgery, prior to starting the recordings, we changed the ventilation of the animal to a mixture of 20% oxygen and 80% medical air. At the same time, we injected a single dose of buprenorphine (0.04 mg/kg, SC; Schering-Plough, UK) and started administering dexmedetomidine that we kept running continuously throughout the recordings (0.075 mg/kg/h, SC; Pfizer Inc., Canada). The isoflurane administration was stopped following the administration of dexmedetomidine and buprenorphine. To examine the effect of anesthesia on the epileptic activity, we performed three additional experiments on SD rats without using ketamine before the intubation. To further examine the effect of anesthesia, we performed seven additional experiments on SD rats anesthetized with a combination of dexmedetomidine (0.075 mg/kg/h, SC) and simultaneous administration of isoflurane (0.3-0.5%), without administering buprenorphine.

The forepaw representation of area S1FL was delineated by optical imaging of the cerebral blood volume (CBV) response to forepaw stimulation. Then, a linear multi-contact probe was inserted into the forepaw region, in an approximately orthogonal orientation relative to the local surface. To determine the effect of dexmedetomidine on seizure susceptibility in a different animal species, we performed similar procedures and the same type of forepaw stimulation in five age-matched mice.

Before the surgery, the mice were injected with the anti-inflammatory drug carprofen (4 mg/kg SC; Zoetis, Canada). The surgical procedure was performed under anesthesia with a solution of ketamine (100 mg/kg IP; Wyeth, Canada) and xylazine (10 mg/kg IP; Bayer Inc., Canada) injected before the surgery, with the addition of isoflurane (0.6-1%; Benson Medical Industries Inc., Canada). The core temperature and heart rate were monitored. The primary somatosensory cortex (S1) of the left hemisphere was exposed, by performing a craniotomy, centered on the forelimb representation in area S1 (S1FL) based on a mouse stereotaxic atlas (AP −0.30 mm, ML ±2.20 mm, DV −1.3 mm; Franklin and Paxinos (2007)). Following the surgery, prior to starting the recordings, the mice were injected with a single dose of buprenorphine (0.1 mg/kg, SC; Schering-Plough, UK). We then started a continuous administration of dexmedetomidine (0.05 mg/kg/h, SC; Pfizer Inc., Canada). Following the administration of dexmedetomidine and buprenorphine, we stopped administering isoflurane.

Then, a linear multi-contact probe was inserted into area S1FL. To maintain the sedation throughout the experiment, we infused dexmedetomidine continuously at a rate of 0.05 mg/kg/h SC. We assessed the depth of sedation by continuously monitoring the vital signs of the animal and by monitoring whether the animal stayed still or performed any movement. In case this monitoring showed that additional sedation was required, we increased the rate of the dexmedetomidine infusion to 0.1 mg/kg/h SC.

### Electrical stimulation of the forepaw

Electrical stimuli were generated using a stimulator/isolator (A365, WPI, Sarasota, FL) and delivered through two needle electrodes inserted into the web spaces of digits 2/3 and 4/5 of the rat or mouse forepaw. With rats and mice, we began our experiments with optical imaging of area S1FL, in order to guide the insertion of the neurophysiology probe to the forepaw representation. Runs for eliciting CBV-based optical imaging responses consisted of ten 4 s long stimulation blocks of 1 ms long, 1 mA electrical current pulses delivered at a frequency of 8 Hz. Following the insertion of the probe, we obtained data for the main experiment, including LFP, HbO, and CBF responses to electrical stimuli delivered to the forepaw. In rats, 8 runs were performed, each consisting of ten 35 s long stimulation trials, separated by ten 55 s long trials in which no stimulus was delivered (see time course in Figure 1A). Each stimulation trial started with 5 s recordings of baseline activity, followed by 10 s of stimulation and 20 s of baseline activity. In thirteen SD rats and five LE rats, each 10 s stimulation block consisted of a train of 1 ms long, 2 mA electrical pulses delivered at a frequency of 8 Hz. In three SD rats, 8 runs were performed with two different stimulus frequencies (4 Hz and 8 Hz) and two different currents (1 mA and 2 mA). In runs 1-2, 3-4, 5-6, and 7-8 we administered 1 mA at 4 Hz, 2 mA at 4 Hz, 1 mA at 8 Hz, and 2 mA at 8 Hz, respectively. In mice, 9 similar runs were performed, except that three different stimulus frequencies (4 Hz, 6 Hz, and 8 Hz) were applied. The recordings were performed in runs with increasing (3 mice) or decreasing (2 mice) stimulation frequency. The duration and intensity of all electrical pulses delivered to mice were 1 ms and 2 mA, respectively. In all experiments in rats and mice, the polarity of stimulation was switched in each pulse relative to the polarity of the preceding pulse.

**Figure 1.**
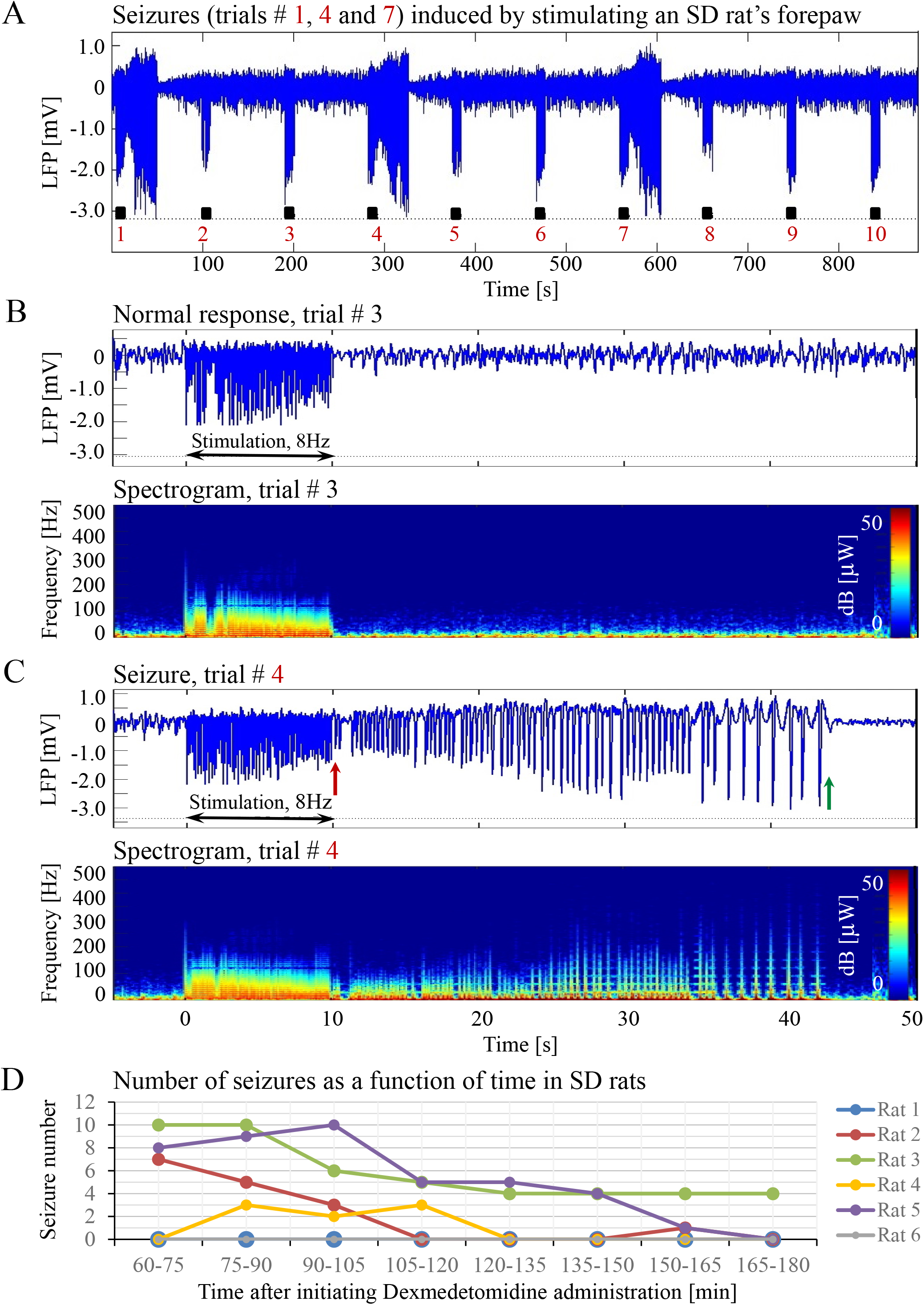
Forepaw stimulation induces epileptic activity in rat area S1FL. **A.** LFP recordings of ten trials, each with 10s-long stimulation. The stimulation periods are marked by black rectangles. Each 10s stimulus consisted of a train of electrical pulses delivered at 8 Hz to the forepaw. Note that during the first, fourth, and seventh trials, the stimulation evoked a seizure. **B.** Top: The LFP (mean averaged over electrode contacts spanning the cortical depth) demonstrates a normal-evoked response in trial #3. Bottom: The corresponding spectrogram (power as a function of frequency and time) computed for the same trial. **C.** Top: LFP (mean averaged over electrode contacts spanning the cortical depth) showing a seizure pattern in trial #4. The red and green arrows indicate the onset and termination, respectively, of a seizure induced by forepaw stimulation. Bottom: The corresponding spectrogram, computed for the same seizure. D. The number of evoked seizures per rat as a function of time shows that seizures were induced already during the first or the second run, only one hour after initiating the dexmedetomidine administration.

### Optical imaging of intrinsic signals

All procedures applied for optical imaging of intrinsic signals (OIS) are described in detail by Bortel et al. (2019). The ROI we imaged was centered on the atlas coordinates of area S1FL in the left hemisphere in rats (AP 0.00 mm, ML ±4.50 mm, DV −2.5 mm; Paxinos and Watson (2005)) and mice (AP −0.30 mm, ML ±2.20 mm, DV –1.3 mm; Franklin and Paxinos (2007)). We imaged the hemodynamic responses to forepaw stimulation at a frame rate of 30 Hz under the illumination of a green and orange LED light with a center wavelength of 530 nm (isosbestic point) and 617 nm, respectively. We computed percent change maps for oxyhemoglobin (HbO) using the modified Beer-Lambert law (Dunn et al., 2003). We applied a correction to adjust for the differential path length through the gray matter at different wavelengths.

### Laser Speckle flowmetry

One laser Speckle diode (Sharp LTO25MD; λ = 798 nm, 30 mW; Thorlabs, Newton, NJ, U.S.A.) was coupled into a 600-μm diameter silica optical fiber (Thorlabs FT600-EMT) with a collimating lens (f = 8 mm, C240-TM; Thorlabs) connected to the distal end of the fiber. The lens was placed approximately 10 cm above the cortical ROI. It was adjusted to provide even illumination over an area with 8 mm diameter on the exposed cortical surface. The coherence length of the laser was approximately 1 cm. The speckle pattern was imaged using a CCD camera (Teledyne Dalsa, Waterloo, Ontario, Canada), which made it possible to obtain 2D maps of CBF at high spatial and temporal resolution. To this end, we quantified the spatial blurring of the speckle pattern that results from blood flow (Boas and Dunn, 2010). Conversion of the raw speckle images to blood flow maps was done using custom-written software that computed the speckle contrast and correlation time values at each pixel (Dunn et al., 2001).

### Electrophysiological data pre-processing and seizure analysis

The methods used for the pre-processing of electrophysiological data and seizure analysis are specified in detail in Bortel et al. (2019).

### Statistical evaluation

We used the Levene’s test to examine whether the variances of two or more compared groups were equal. In the cases where equal variances were verified, a Student’s t-test was performed to test a difference between the means of two groups. If there was evidence to support non-equal variances, a Mann-Whitney or post hoc Tamhane’s test was applied to evaluate statistical differences between two or more groups, respectively. A nonparametric Wilcoxon test was applied to groups of paired data variables. Results are presented as a mean ± SEM. Differences with p < 0.05 were considered statistically significant (IBM SPSS Statistics, 2016).

## RESULTS

### Characterization of normal evoked responses and epileptic responses

Typical LFP patterns recorded during normal evoked responses in area S1FL of an SD rat are presented in Figures 1A (trials 2, 3, 5, 6, 8, 9, and 10) and 1B. The normal evoked responses were confined within the 10 s period of stimulation. As illustrated in Figure 1B, the LFP amplitudes in response to electrical stimulation pulses were higher than the spontaneous LFP amplitudes. In four of six rats, we detected seizures induced by somatosensory stimulation of the forepaw. Figures 1A (trials 1, 4, and 7) and 1C present seizures induced by the forepaw stimulation. As can be observed in Figure 1C, the onset of the seizure (marked by a red arrow) consisted of high-frequency negative or positive-going deflections of the mean extracellular field potential.

The seizures typically extended for several seconds longer than the stimulation period. Before they terminated, the seizures consisted of low-frequency, high amplitude deflections of the field potential, extending above or below the baseline (Figure 1C; the seizure end is marked by a green arrow). The LFP and the corresponding spectrograms were used to estimate the seizure onset and termination times, relative to the first stimulation pulse. All seizures evoked in our rats were brief, lasting less than one minute. The average duration of seizures was 31.50 ± 1.83 s. The seizures were followed by another seizure (55 cases out of 113 seizures), or a refractory period (40 refractory periods out of 113 seizures) or a normal response (14 normal responses out of 113 seizures).

The first recording run was performed 15 min after the isoflurane administration was stopped, thus 30 min following the administration of dexmedetomidine and buprenorphine. In the first 10-min run, we stimulated the forepaw for eliciting CBV-based optical imaging responses (4 s stimulation blocks, administering 1 mA pulses at 8 Hz). We did not observe any epileptic activity during this first stimulation run. Approximately 40 min following the administration of dexmedetomidine and buprenorphine, prior to the stimulation runs, we acquired two 10 min long runs of spontaneous activity. We did not observe any epileptic activity during these runs. These findings indicate that dexmedetomidine alone does not induce seizures: for generating seizures, potent stimuli are required too.

Seizures were observed already after one hour of dexmedetomidine administration, during the first or the second run in which we applied the potent stimulation: 10 s stimulation blocks of 2 mA pulses at 8 Hz (Figure 1D). In SD rats, the average number of observed seizures during the period of 60-120 min following the initiation of dexmedetomidine administration (14.3±5.9, median equal 11.5) was larger than the corresponding number observed during the period of 120-180 min following this initiation (4.5±2.8, median equal 0.5, n = 6 rats; p=0.07, Wilcoxon test; non-significant trend).

The seizures induced in rats by electrical forepaw stimulation were exclusively electrographic seizures. We did not observe any epileptic behavior.

### Seizures induced by forepaw stimulation under dexmedetomidine sedation are associated with extended cerebral blood oxygenation and flow responses

To test how the seizures induced by forepaw stimulation under dexmedetomidine sedation influence hemodynamic responses, we analyzed the spatial extent of the hemodynamic responses elicited during normal evoked responses and seizures. Figures 2A-B and 3A-B present the spatial HbO and CBF responses, respectively, 1 s before the stimulus onset (1), 10 s after the stimulus onset (2), and 10 s after the cessation of the stimulus (3), for normal evoked response and seizure recorded in a single trial in one animal.

**Figure 2.**
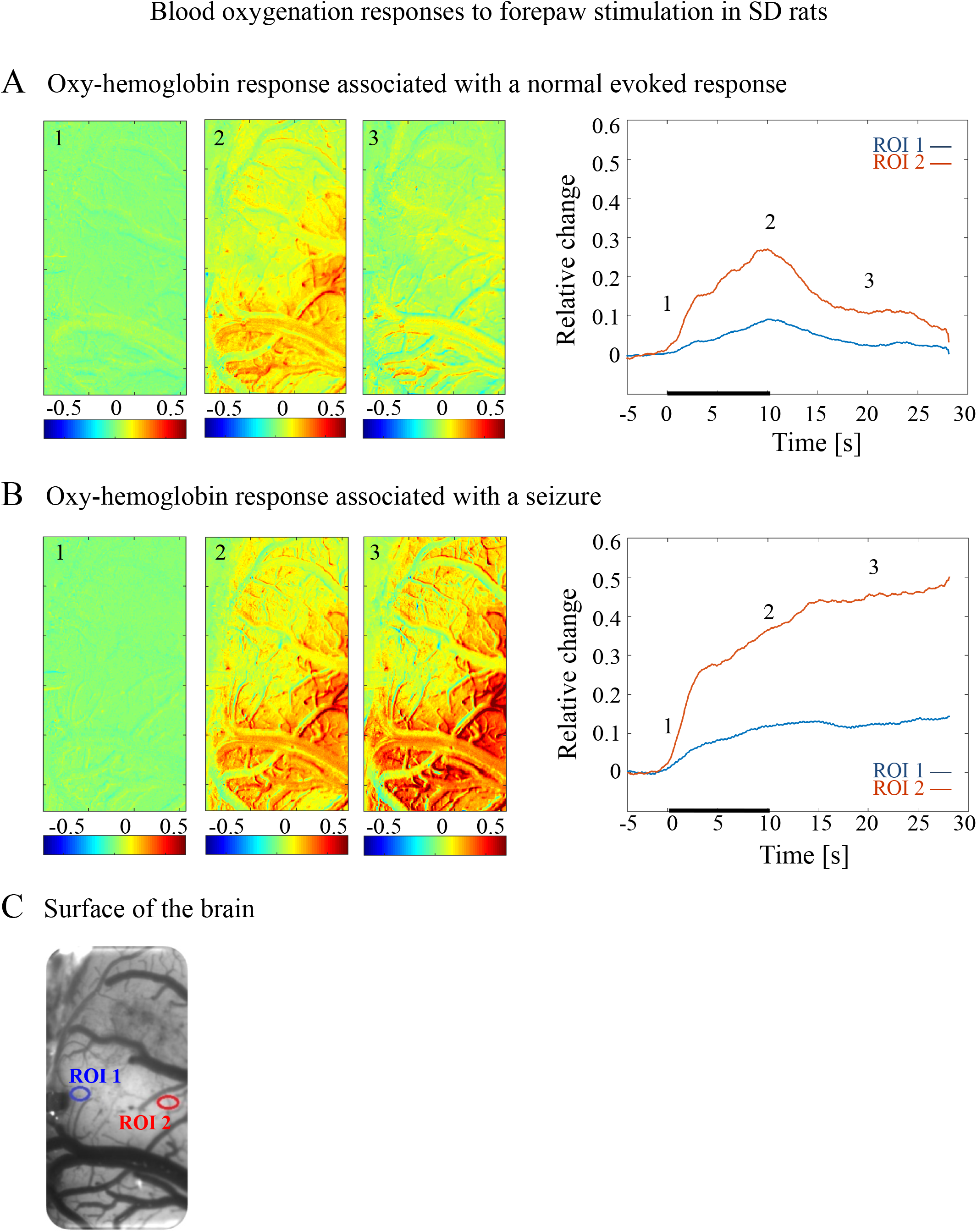
Cerebral blood oxygenation responses evoked by forepaw stimulation. **A.** Cerebral blood oxygenation response evoked by stimulation of the contralateral forepaw. To the left, the spatial responses before (the 1s before stimulus onset), during (9-10s following stimulus onset), and after (9-10s following the cessation of the stimulus) the 10s-long forepaw stimulation period. The reference for obtaining these responses was imaged between 3 and 1 seconds before stimulus onset. Note that positive response indicated in indexed yellow and red colors represent an increase in blood oxygenation. To the right are two time-courses presenting the corresponding temporal response from two regions (blue and red ROIs in panel C) within the activated area. The stimulation period between 0 and 10 seconds is marked by a dark bar. **B.** Maps of the blood oxygenation changes during a response that evoked a seizure, from before, during, and after the 10s-long forepaw stimulation period (exact time periods are as in A). To the right are two time-courses presenting the corresponding temporal response from two regions (blue and red ROIs in panel C) within the activated area. **C.** The imaged cortical surface with the two ROIs used for sampling the time-courses presented in A and B.

**Figure 3.**
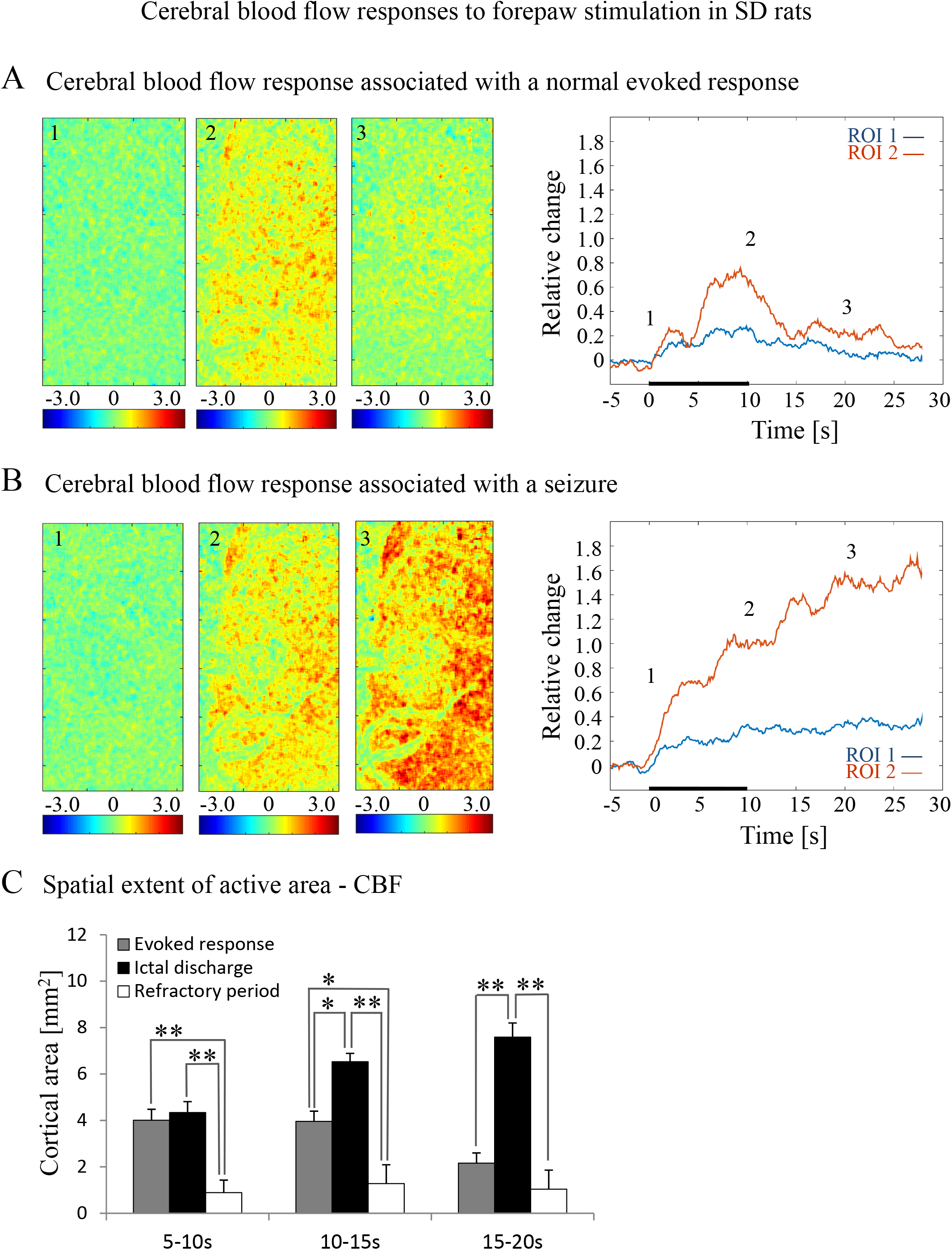
Cerebral blood flow responses evoked by forepaw stimulation. **A.** Cerebral blood flow response evoked by stimulation of the contralateral forepaw. To the left, the spatial responses before (the 1s before stimulus onset), during (9-10s following stimulus onset), and after (9-10s following the cessation of the stimulus) the 10s-long forepaw stimulation period. The reference for obtaining these responses was imaged between 3 and 1 seconds before stimulus onset. Note that positive responses indicated in indexed yellow and red colors represent an increase in blood flow. To the right are two time-courses presenting the corresponding temporal response from two regions (blue and red ROIs in Figure 2C) within the activated area. The stimulation period between 0 and 10 seconds is marked by a dark bar. **B.** Maps of the blood flow changes during a response that evoked a seizure, from before, during, and after the 10s-long forepaw stimulation period (exact time periods are as in A). To the right are two time-courses presenting the corresponding temporal response from the same ROIs as described in A. **C.** A bar graph showing the spatial extent of the CBF response calculated for the epochs of 5-10s, 10-15s, and 15-20s relative to the onset of the stimulus during normal evoked responses (n=21), seizure responses (n=20) and refractory periods (n=14; * p<0.05, ** p<0.001; Tamhane’s test).

The spatial extent of the cerebrovascular hemodynamic responses 10 s following the onset of stimulation was slightly larger during a seizure (Figure 2B and 3B, panel 2) than during a normal response (Figure 2A and 3A, panel 2). The spatial extent of the responding region 10 s following the cessation of the stimulus was substantially larger during a seizure (Figure 2A and 3A, panel 3) than during a normal response (Figure 2B and 3B, panel 3). The time-courses of the HbO and CBF changes during a normal evoked response and an induced seizure response are presented on the right panels of Figures 2A-B and 3A-B, respectively. The two different colored lines correspond to the two brain regions marked in Figures 2C. At any time-point following the onset of the stimulus, the HbO and CBF amplitudes during the seizure (Figure 2B and 3B) were higher than the corresponding amplitudes during the normal evoked response (Figure 2A and 3A). The bar plot in Figure 3C presents the spatial extent of the CBF responses averaged over six animals, during normal responses, seizures, and refractory periods, respectively. During the period of 5–10 s relative to the onset of the stimulus, the spatial extent of the cerebrovascular hemodynamic responses during seizures was larger than during normal refractory periods (Figure 3C; ** p < 0.001, Tamhane’s test). For each of the periods of 10–15 s, and 15–20 s relative to the onset of the stimulus, the spatial extent of the cerebrovascular hemodynamic responses during seizures was larger than during normal evoked responses and refractory periods (Figure 3C; * p < 0.05, ** p < 0.001, Tamhane’s test). Note that these two periods – namely 10–15 s and 15–20 s relative to the onset of the stimulus – are part of the post-stimulation period in which the seizures persist with no sensory stimulation. Interestingly, not only the average spatial extent of the seizures was larger than that of normal responses obtained during the same period following the cessation of the stimulus, also the seizure’s spatial extent *post-stimulation* was larger than that observed in normal responses *during stimulation* (Figures 3C; p < 0.001, Tamhane’s test). Thus, the hemodynamic responses elicited by the seizures propagated beyond the spatial extent of the normal responses to forepaw stimulation.

### The increased susceptibility to seizures does not depend on using ketamine, buprenorphine, or combining isoflurane with dexmedetomidine

To evaluate the dependence of seizure generation on the anesthesia regime, we first performed 3 experiments under dexmedetomidine and buprenorphine sedation but without using the anesthetic ketamine before the intubation and surgery. All three rats had seizures in response to forepaw stimulation of 8 Hz and 2 mA (Figure 4 A-D).

**Figure 4.**
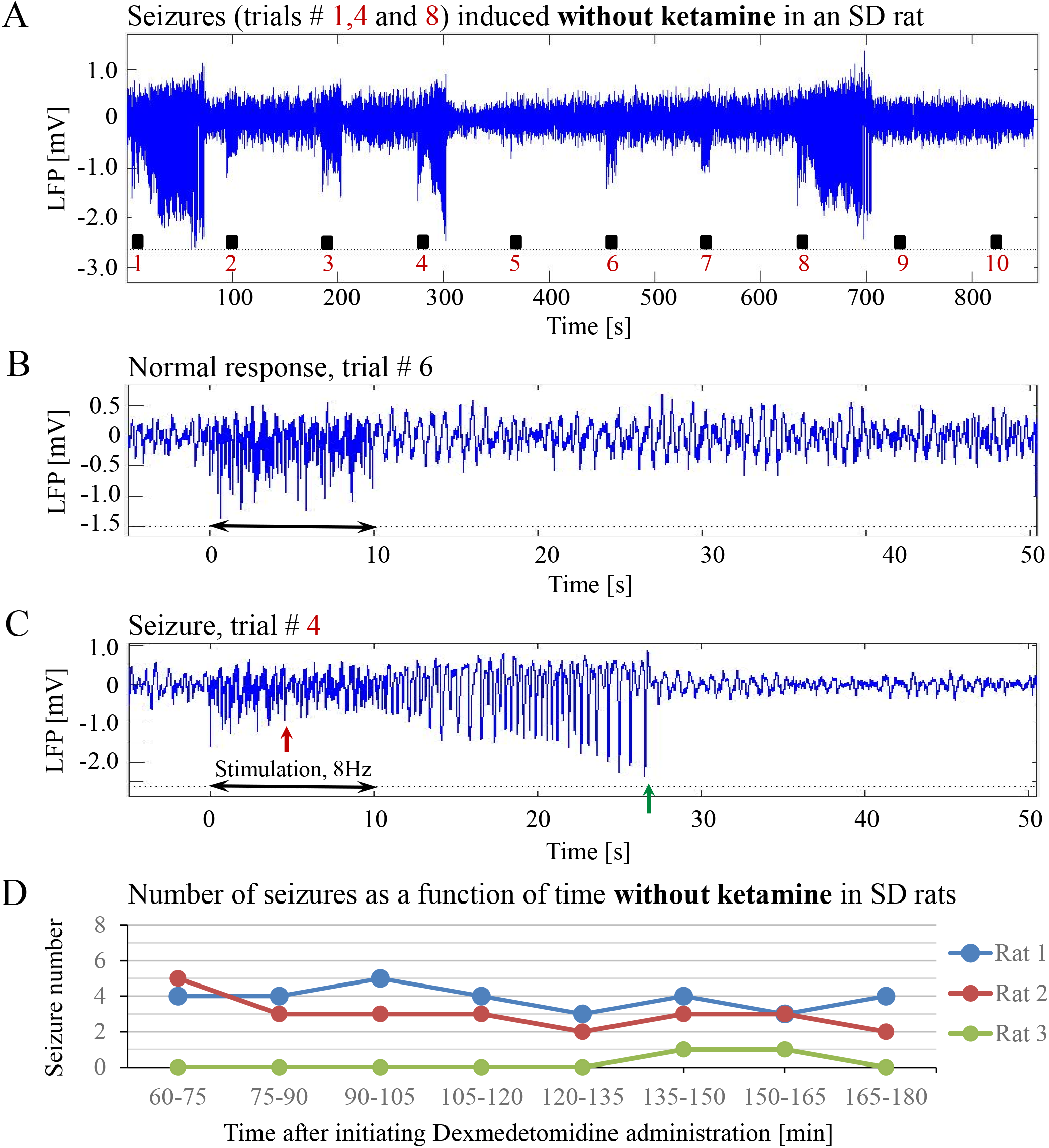
The increased susceptibility to seizures in SD rats under dexmedetomidine and buprenorphine does not depend on ketamine. The data presented in this figure were obtained from 3 rats under dexmedetomidine and buprenorphine sedation with no administration of ketamine. The format of the presentation is similar to that used in Figure 1. **A.** A time-course of LFP (averaged over electrode contacts spanning the cortical depth) recorded during ten trials, each with 10s-long stimulation. The stimulation periods are marked by black rectangles. Note the seizures induced by the stimulus in the first, fourth, and eighth trials. **B.** A normal-evoked LFP response recorded in trial #6. **C.** A seizure pattern recorded in trial #4. The red and green arrows indicate the onset and termination of the seizure, respectively. **D.** The number of seizures recorded in each of the 3 rats as a function of time after the initiation of the dexmedetomidine administration.

Previous studies reported that isoflurane protects against seizures and that combining dexmedetomidine with isoflurane anesthesia prevents seizures in response to forepaw stimulation. To test these assertions, we performed four additional experiments using SD rats anesthetized with a combination of dexmedetomidine and 0.3%-0.5% isoflurane, without administering buprenorphine. In response to forepaw stimuli of 2 mA administered at 8 Hz, we observed seizures in 50% of these rats (2 out of 4; Figure 5A-D). We concluded that ketamine and buprenorphine are not necessary for the effect of increasing susceptibility to seizures. We also concluded that combining isoflurane with dexmedetomidine does not eliminate the effect of increased susceptibility to seizures.

**Figure 5.**
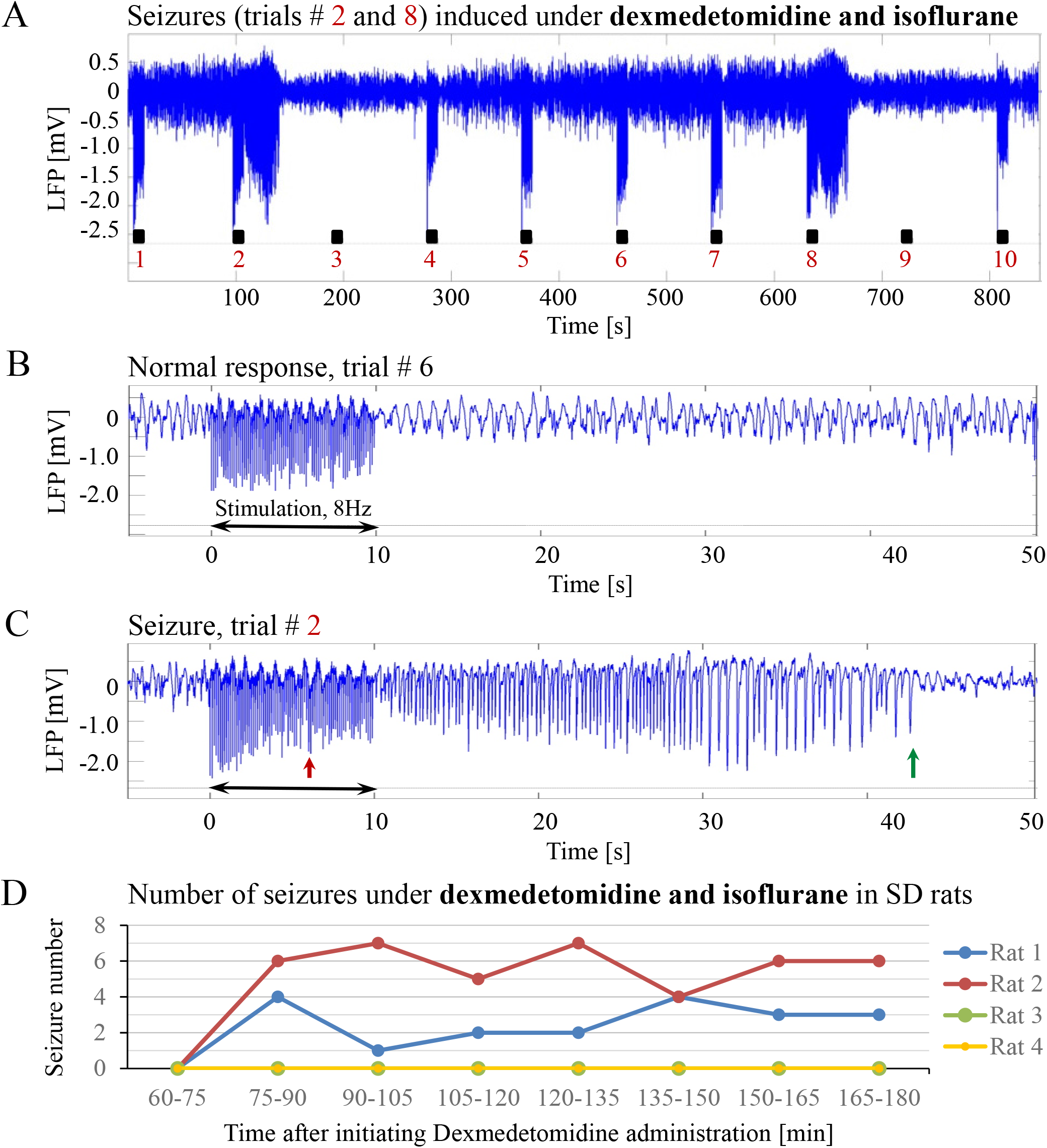
Potent forepaw stimulation induces seizures in SD rats under dexmedetomidine and isoflurane anesthesia. The data presented in this figure were obtained from 4 rats under dexmedetomidine and isoflurane anesthesia. **A.** LFP recordings of ten trials, each with 10s-long stimulation. Note the seizures induced by the stimulus in the second and eighth trials. **B.** A normal-evoked LFP response recorded in trial #6. **C.** A seizure pattern recorded in trial #2. **D.** The number of seizures recorded in each of the 4 rats as a function of time after the initiation of the dexmedetomidine administration. Seizures were induced already during the second run, less than 90 minutes after initiating the dexmedetomidine administration.

### The susceptibility to seizures depends on the stimulation paradigm: seizures-free stimulation parameters

Our results show that under dexmedetomidine sedation, forepaw stimulation with pulses of 2 mA administered at 8 Hz generates seizures. To test the dependence of the seizure generation on the electrical stimulation parameters, we administered forepaw stimuli with different currents (1 mA and 2 mA) and frequencies (4 Hz and 8 Hz) in 3 SD rats under dexmedetomidine and buprenorphine sedation. Forepaw stimuli of 1 mA and 2 mA administered at 4 Hz did not generate seizures (Figure 6A-B). In contrast, administering electrical pulses of 1 mA and 2 mA at 8 Hz induced seizures in 2 out of 3 and all 3 rats, respectively (Figure 7A-C). Decreasing the stimulation current and frequency made it possible to obtain seizure-free response to forepaw stimulation under dexmedetomidine and buprenorphine sedation (Fig. 6). This indicates that similar decreases may result in seizure-free forepaw responses under dexmedetomidine and isoflurane too. We, therefore, aimed to find a seizure-free forepaw stimulation paradigm under dexmedetomidine and isoflurane sedation. To this end, we performed experiments in 3 additional SD rats whose forepaw was stimulated with 1.5 mA electrical pulses administered at 4 Hz. Under this condition, we did not observe any seizure (Figure 8).

**Figure 6.**
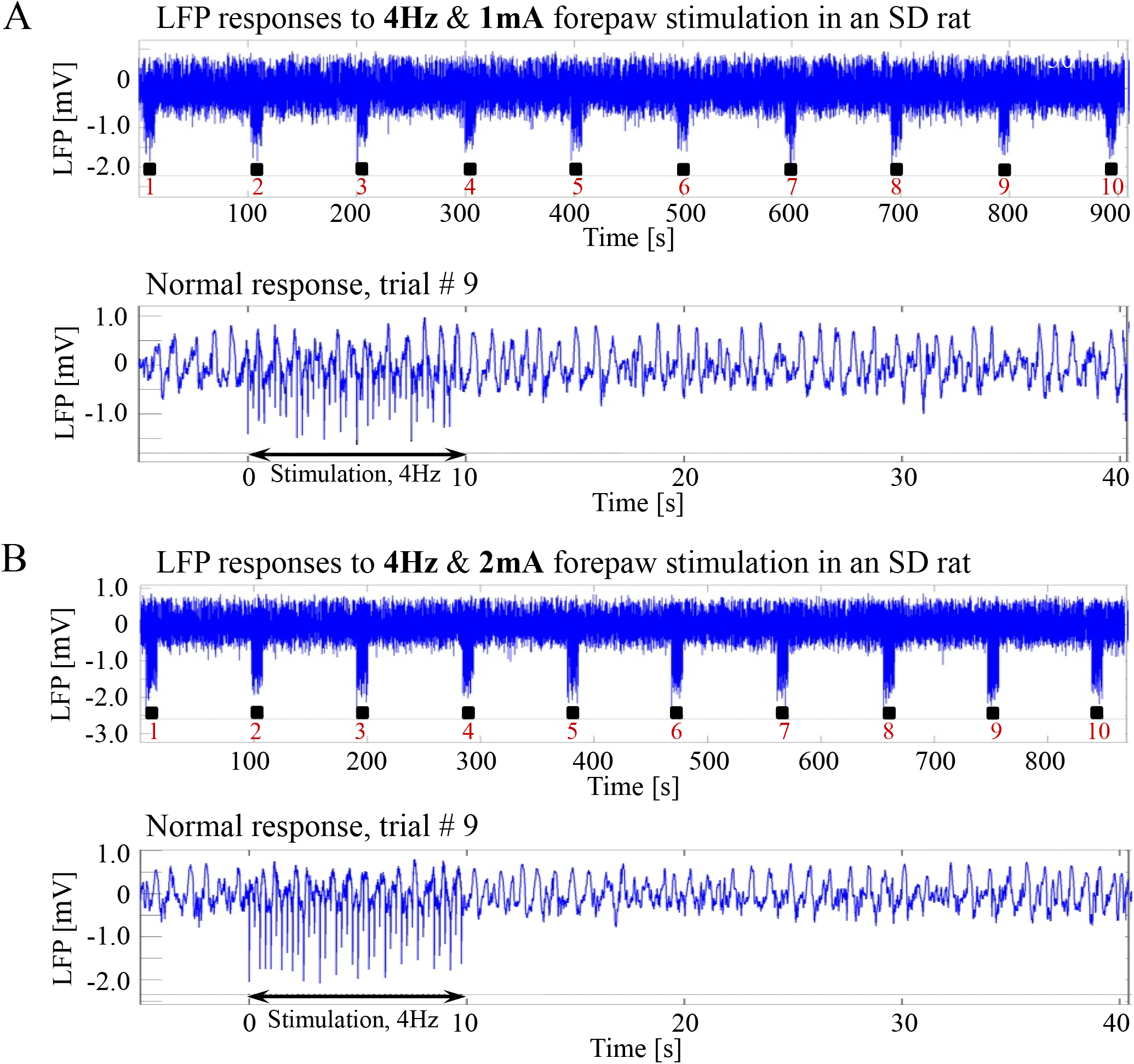
Normal LFP responses to forepaw stimulation administered at 4Hz in SD rats under dexmedetomidine and buprenorphine. The data presented in this figure were obtained from 3 rats under dexmedetomidine and buprenorphine sedation. **A.** Top: Normal LFP responses obtained from area S1FL, in response to **4Hz & 1mA** forepaw stimulation. The panel shows ten trials, each with 10s-long stimulation. Bottom: A magnification of the normal evoked response from the ninth trial. **B.** Top: Normal LFP responses to **4Hz & 2mA** forepaw stimulation. Bottom: A magnification of the normal evoked response from the ninth trial.

**Figure 7.**
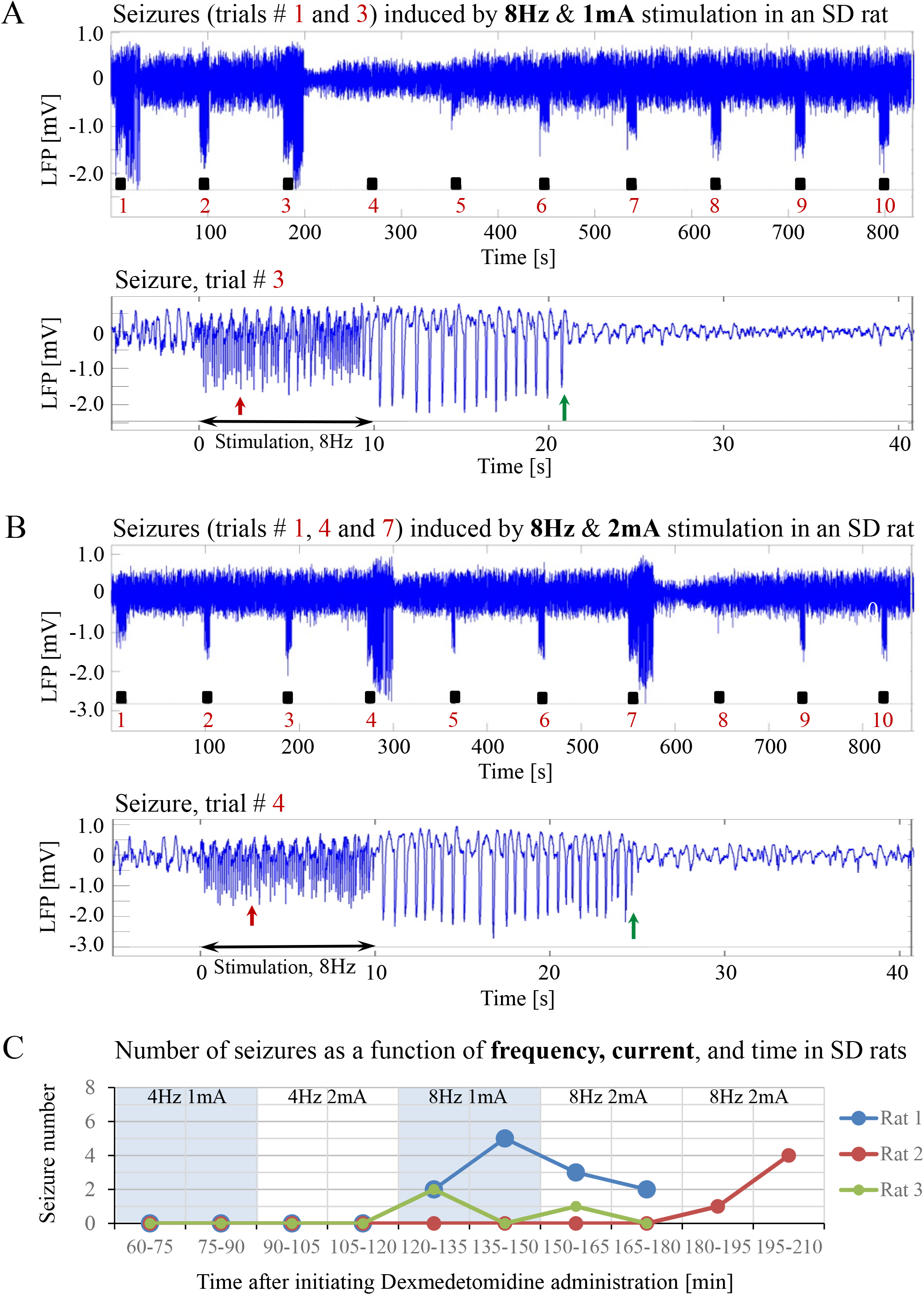
Forepaw stimulation administered at 8Hz induces epileptic activity in SD rats under dexmedetomidine and buprenorphine. The data presented in this figure were obtained from the same 3 rats whose responses to forepaw stimulation with different parameters are presented in Figure 6. **A.** Top: LFP responses to **8Hz & 1mA** forepaw stimulation. Note the seizures induced by the stimulus in the first and third trials. Bottom: A magnified view of the seizure recorded in the third trial. **B.** Top: LFP responses to **8Hz & 2mA** forepaw stimulation. Note the seizures induced by the stimulus in the first, fourth, and seventh trials. Bottom: A magnified view of the seizure recorded in the fourth trial. **C.** The number of seizures recorded in each of the 3 rats whose responses to forepaw stimulation are presented in Figures 6 and 7. The number of seizures is presented as a function of time after the initiation of the dexmedetomidine administration. The stimulation parameters used for each pair of consecutive blocks are presented at the upper part of the panel.

**Figure 8.**
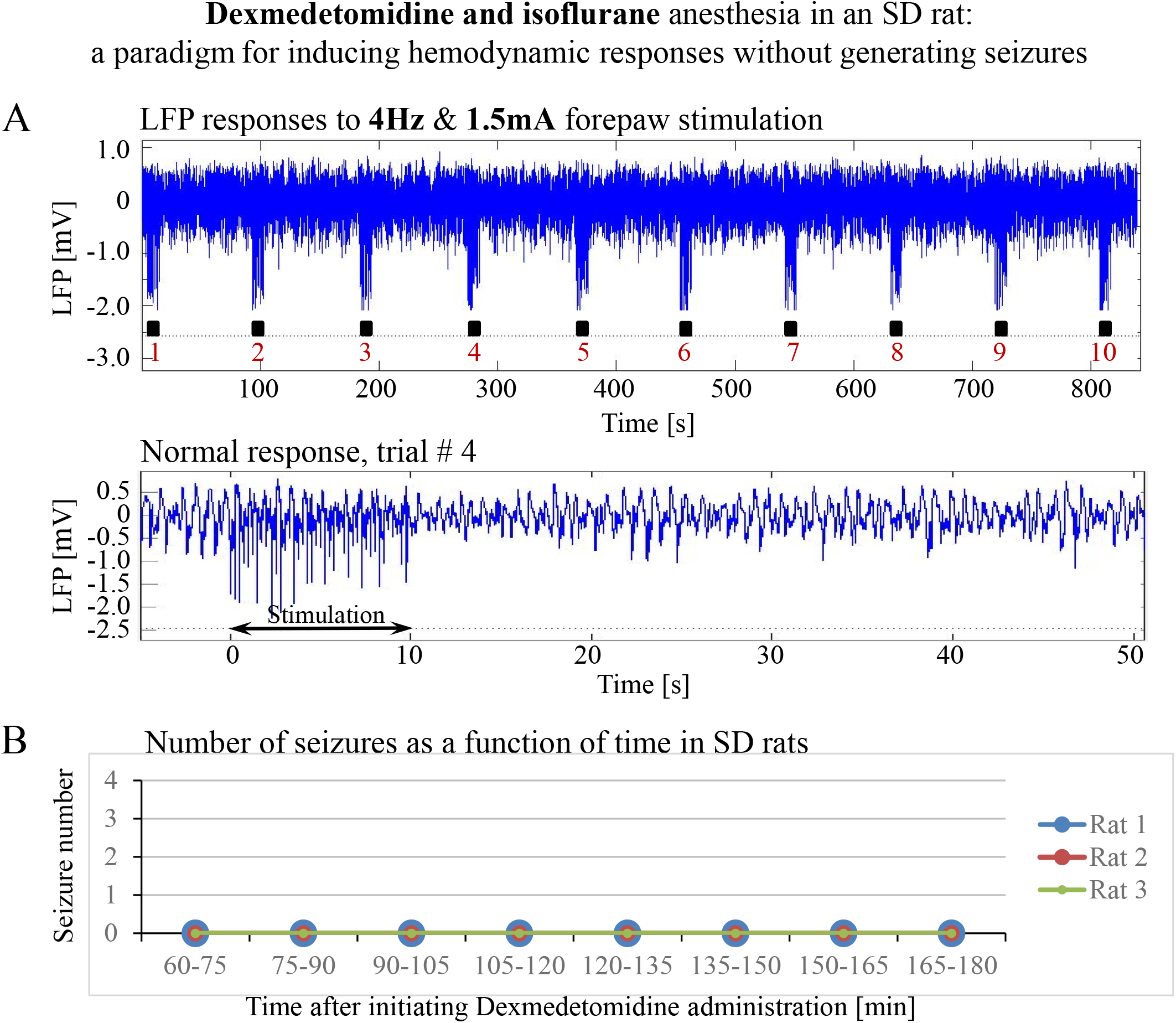
A paradigm for seizure-free forepaw stimulation under dexmedetomidine and isoflurane. The data presented in this figure were obtained from 3 rats under dexmedetomidine and isoflurane anesthesia. Forepaw stimuli of 1.5 mA were administered at a rate of 4Hz. **A.** Top: Normal LFP responses to 10s-long stimulation blocks obtained from area S1FL. Bottom: A magnification of the normal evoked response from the fourth trial. **B.** The number of seizures (none) recorded in each of the 3 rats as a function of time shows that forepaw stimuli of 1.5mA administered at 4Hz do not induce seizures in SD rats under dexmedetomidine and isoflurane anesthesia.

### Hemodynamic responses associated with the seizure-free forepaw stimulation paradigms

To compare and evaluate the detectability of the hemodynamic responses associated with the three seizure-free paradigms we have demonstrated (Figure 6A-B and Figure 8), we first analyzed the CBV responses obtained in single-trials under these paradigms. Figures 9A and 9B compare the spatial extent and time courses obtained under dexmedetomidine and buprenorphine in response to stimulation at 4 Hz with 1 mA and 2 mA pulses, respectively. Figure 9C presents the spatial extent and time courses obtained under dexmedetomidine and isoflurane in response to 1.5 mA pulses administered at 4 Hz. All three paradigms generated single-trial hemodynamic responses that were well detectable.

**Figure 9.**
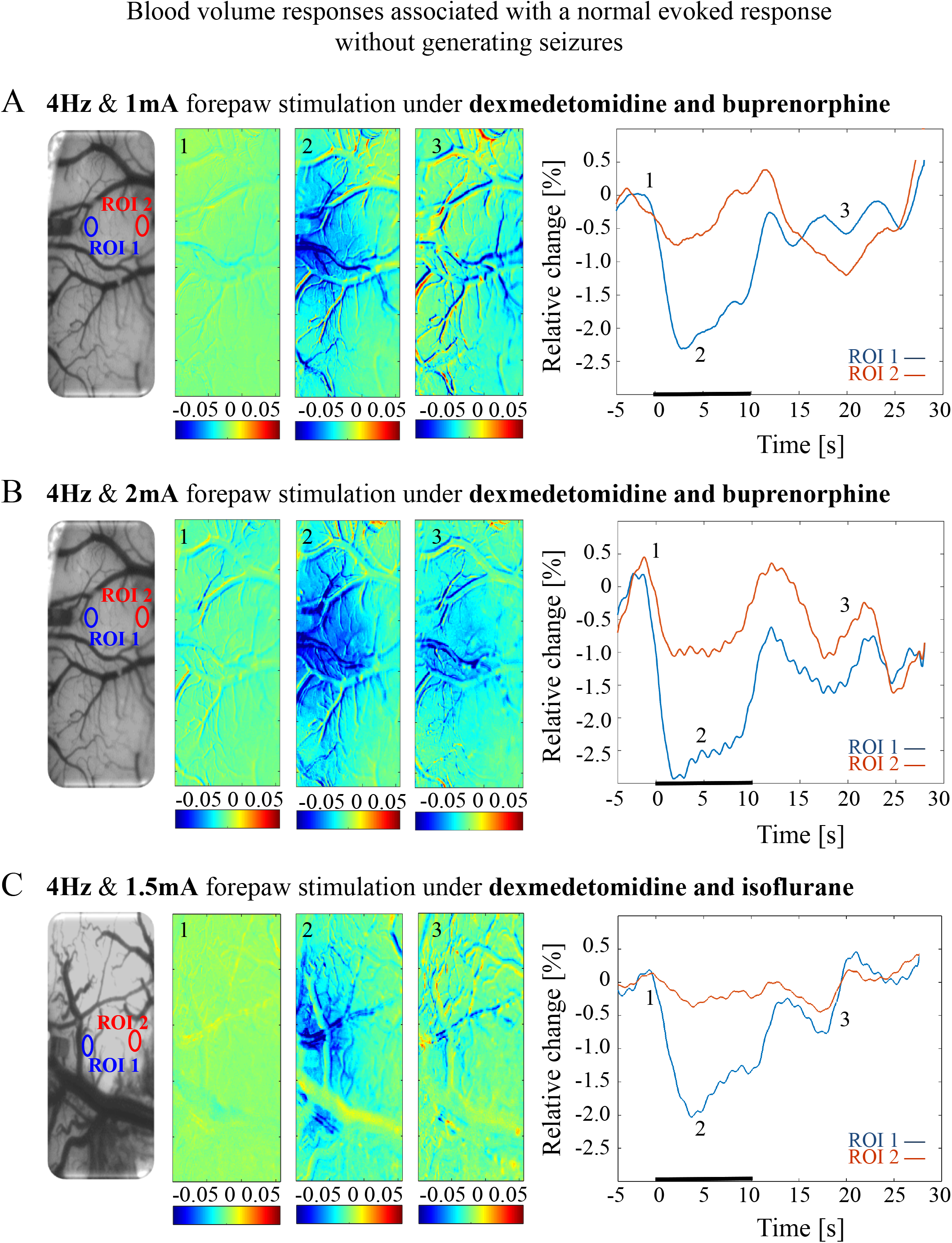
Cerebral blood volume responses associated with seizure-free responses to 4Hz forepaw stimulation under different anesthesia regimes. **A.** A cerebral blood volume response to a 10s-long block of forepaw stimuli of **1mA administered at 4Hz** under **dexmedetomidine and buprenorphine** sedation. To the left, the imaged cortical surface. The 3 maps presented in color show the spatial responses before (the 1s before stimulus onset), during (4-5s following stimulus onset), and after (9-10s following the cessation of the stimulus) the 10s-long forepaw stimulation period. The reference for obtaining these responses was imaged between 3 and 1 seconds before stimulus onset. Note that negative responses indicated in indexed blue colors represent an increase in blood volume. To the right are two time-courses presenting the corresponding temporal response from two regions (blue and red ROIs in the left panel) within the activated area. The stimulation period between 0 and 10 seconds is marked by a dark bar. **B.** Cerebral blood volume response evoked by **4Hz and 2mA** forepaw stimulation under **dexmedetomidine and buprenorphine** sedation. The format of the presentation is identical to that used in panel A. **C.** Cerebral blood volume response evoked by **4Hz and 1.5 A** forepaw stimulation under **dexmedetomidine and isoflurane** anesthesia. The format of the presentation is identical to that used in panel A.

The CBV responses obtained under dexmedetomidine and buprenorphine were higher in response to 2 mA pulses than to 1 mA pulses (p<0.001, Tamhane’s test; Figure 10). They were also higher than the responses to 1.5 mA pulses obtained under dexmedetomidine and isoflurane sedation (p<0.001; Figure 10).

**Figure 10.**
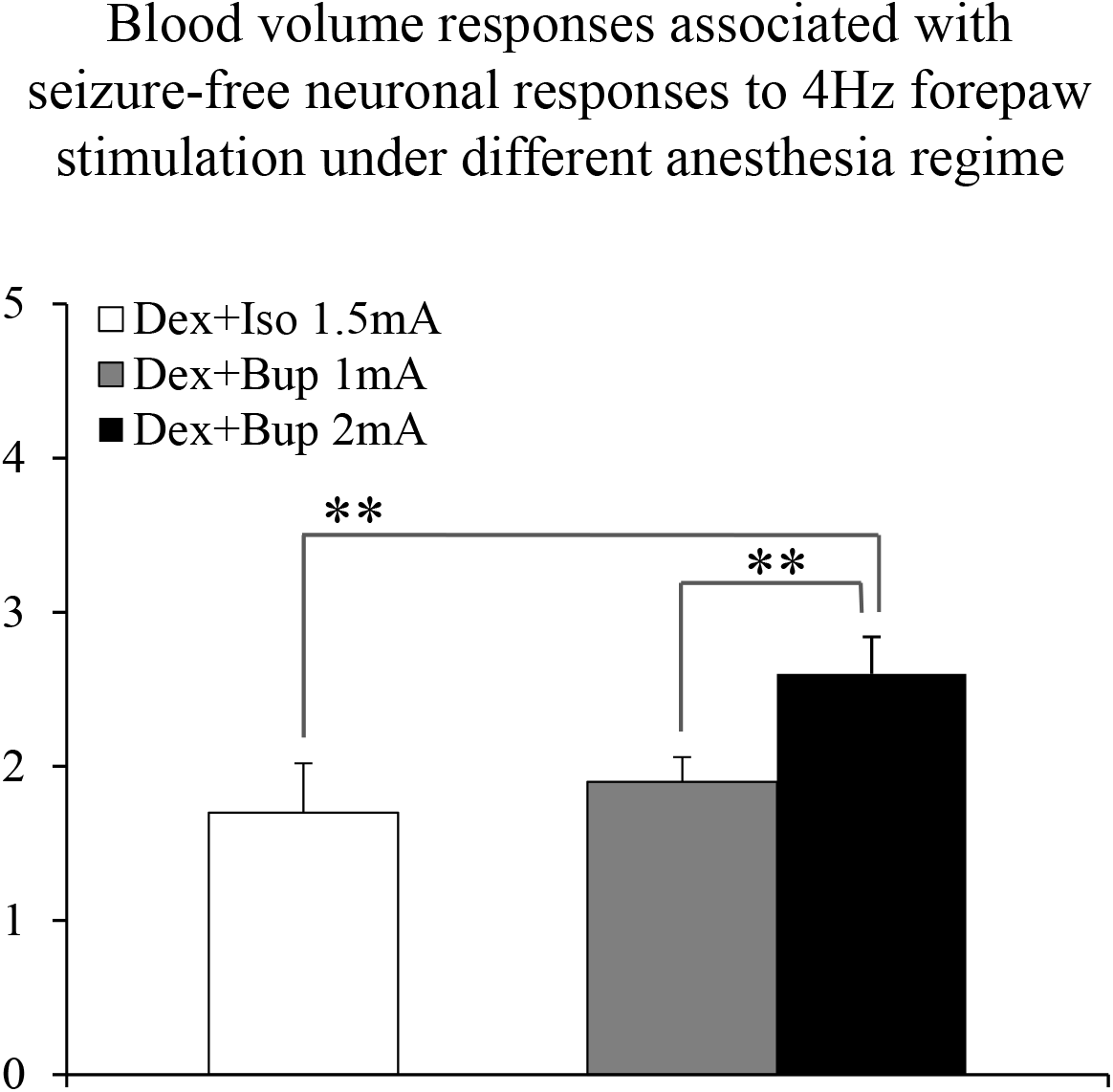
The amplitudes of cerebral blood volume responses associated with seizure-free responses to 4Hz forepaw stimulation under different anesthesia regimes. A bar graph showing the peak amplitude of CBV normal responses sampled from the gray matter region that showed the highest response. The white bar to the left shows the amplitude of response to stimuli of 1.5mA administered at 4Hz under dexmedetomidine and isoflurane anesthesia (Dex + Iso; mean ± SEM, n=60 trials from 3 SD rats). The gray and black bars show the amplitudes of responses to stimuli of 1.0mA and 2.0mA, respectively, administered at 4Hz under dexmedetomidine and buprenorphine sedation (Dex + Bup; n=60 trials from the same 3 SD rats). ** marks p<0.001; Tamhane’s test.

### Long Evans rats under dexmedetomidine sedation are susceptible to seizures; wild type mice are not

The susceptibility of the brain to seizure generation is known to depend on animal species (Schauwecker, 2002). In addition to SD rats, Long-Evans (LE) rats and mice of different strains are commonly used for studying the resting-state and neurovascular coupling. To test the susceptibility of these strains to seizure generation under dexmedetomidine sedation, we pursued similar experiments using LE rats and C57BL6 wild type mice.

Forepaw stimulation with current of 2 mA administered at 8 Hz elicited seizures in all 5 LE rats we tested under dexmedetomidine and buprenorphine sedation. The LFP characteristics of seizures evoked in LE rats were similar to those evoked in SD rats (Figure 11A-D). Seizures induced in LE rats were observed as soon as one hour after the dexmedetomidine administration, during the first or second stimulation run (Figure 11D). The average number of seizures during the period of 60-120 min following the initiation of dexmedetomidine administration (15.8±5.6, median equal 17) was not statistically different than the corresponding number observed during the period of 120-180 min following this initiation (14.0±5.2, median equal 20; p=0.34 Wilcoxon test). The seizures were followed by another seizure (72 cases out of 149 seizures), or a normal response (60 normal responses out of 149 seizures), or a refractory period (16 refractory periods out of 149 seizures; Figure 11A-C). Lastly, 100% of LE rats had seizures, in comparison to 67% of the SD rats from which we obtained data from 8 runs. We observed a non-statistically significant trend of a higher number of seizures in LE rats (29.8 ± 10.7 per animal) than the corresponding number observed in SD rats (18.8 ± 8.5 per animal; p=0.43, Mann-Whitney’s test).

**Figure 11.**
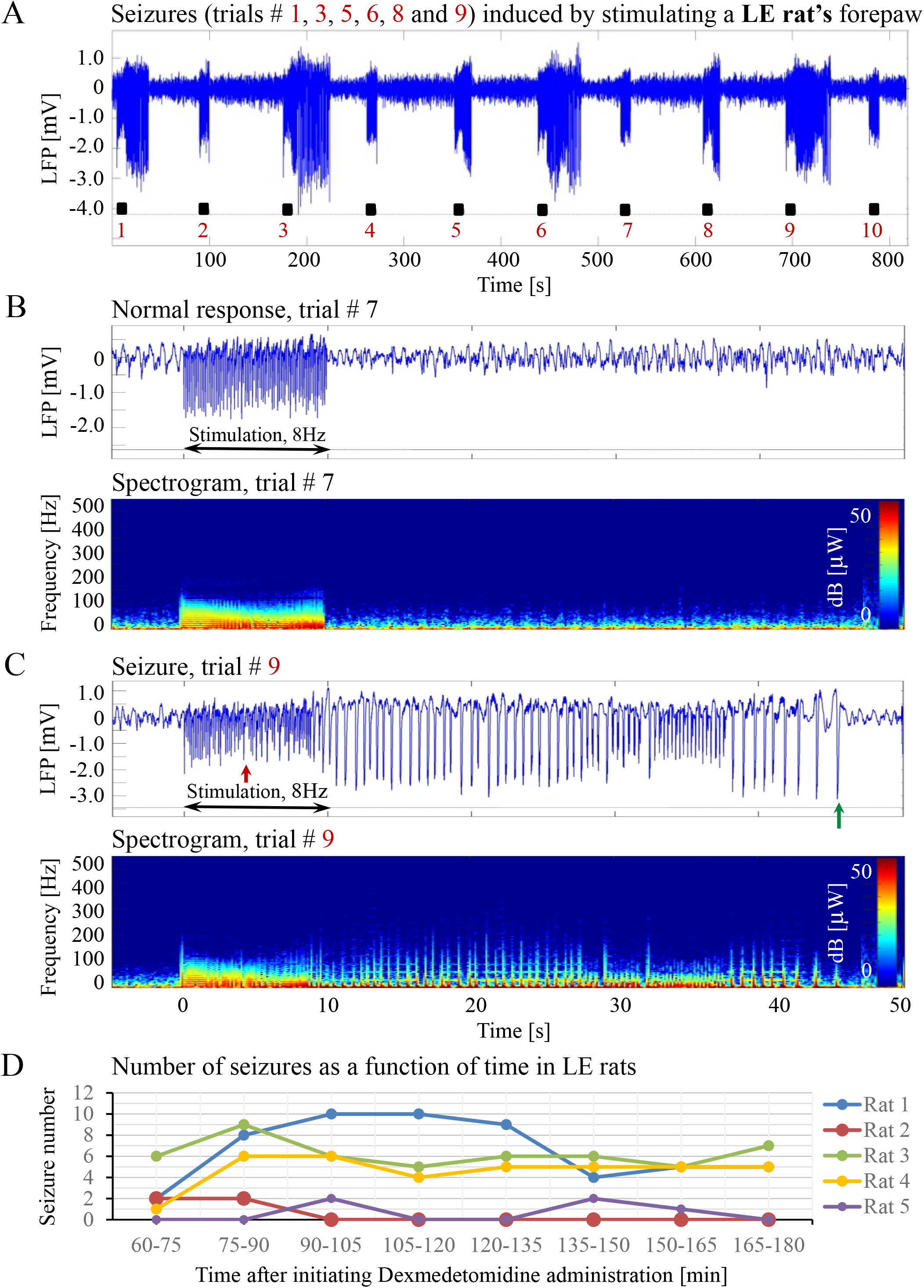
Forepaw stimulation induces epileptic activity in LE rat. **A.** LFP recordings of ten trials, each with 10s-long stimulation. The stimulation periods are marked by black rectangles. Note that during the first, third, fifth, sixth, eight, and ninth trials, the stimulation induced a seizure. **B.** Top: The LFP (mean averaged over electrode contacts spanning the cortical depth) demonstrates a normal-evoked response in trial #7. Bottom: The corresponding spectrogram (power as a function of frequency and time) computed for the same trial. **C.** Top: LFP (mean averaged over electrode contacts spanning the cortical depth) showing a seizure pattern in trial #9. The red and green arrows indicate the onset and termination, respectively, of a seizure induced by forepaw stimulation. Bottom: The corresponding spectrogram, computed for the same seizure. D. The number of evoked seizures per rat as a function of time shows that seizures are induced already during the first or the third run, only one hour after initiating the dexmedetomidine administration.

In pilot experiments in 2 mice, we did not observe seizures and not even high-amplitude LFP responses to stimuli of 2 mA administered at 8 Hz, identical to those that elicited high-amplitude responses in SD and LE rats. We hypothesized that the reason for the lack of high-amplitude responses was that the inhibition elicited by the response in mice remains effective for a longer duration than it does in rats. In addition, lower frequency 6 Hz, 3 s corneal stimulation previously produced ‘psychomotor’ seizures in mice (Barton et al., 2001; Esneault et al., 2017). Based on these observations, we decided to administer forepaw stimuli to five age-matched mice using three different frequencies of stimulation: 4 Hz, 6 Hz, and 8 Hz. Figures 12A and B present normal responses obtained with 4 Hz and 6 Hz stimulation, respectively. Note that the normal responses to stimuli are noticeable with the 4 Hz – the lowest stimulation frequency – and are substantially weaker with 6 Hz stimulation. The LFP responses to forepaw stimuli administered at 8 Hz were virtually undetectable (Figure 12C). As illustrated in Figure 12A-C, we did not evoke any seizure with any of the stimulation frequencies we administered to any of the mice under dexmedetomidine and buprenorphine. In conclusion, seizures were observed in SD and LE rats, but not in C57 mice.

**Figure 12.**
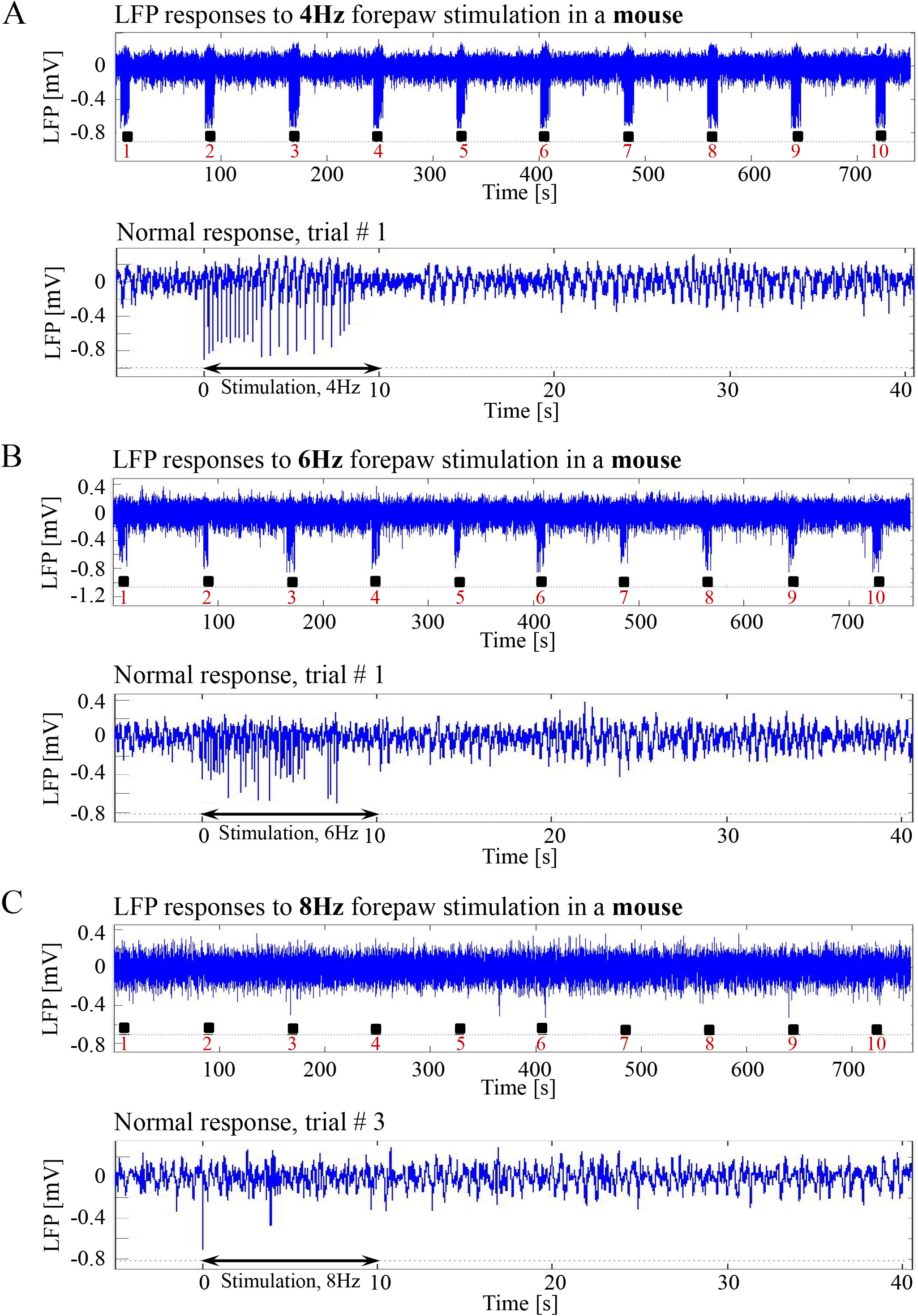
Absence of seizures in the C57BL6 mouse strain. **A.** Top: Normal LFP responses obtained from area S1FL, in response to 4Hz forepaw stimulation. The panel shows ten trials, each with 10s-long stimulation. The stimulation blocks are marked by black rectangles. Bottom: A magnification of the normal evoked response from the first trial. **B.** Top: Normal LFP responses to 6Hz forepaw stimulation. Bottom: A magnification of the normal evoked response from the first trial. **C.** Top: A typical activity pattern of evoked responses to 8Hz forepaw stimulation. Bottom: A magnification of the normal evoked response from the third trial.

## DISCUSSION

### Dexmedetomidine sedation increases susceptibility to seizures

We have demonstrated that under sedation or anesthesia that involve dexmedetomidine, long (10 s), potent (1 mA or 2 mA), repetitive and frequent (8 Hz) peripheral electrical stimuli evoke seizures (Figures 1-5 and 7A-C). Epileptic activity induced in our rats consisted of brief and focal electrographic seizures; the animals did not show any epileptic behavior.

To identify the compound that causes the increased susceptibility to seizures, we managed several sedation and anesthetic regimes by administering different combinations of the following compounds: ketamine, xylazine, carprofen, dexmedetomidine, buprenorphine, isoflurane, and urethane (we used the latter in Bortel et al., 2019).

To test whether ketamine could be the compound that increases the susceptibility to seizures, we performed three experiments under dexmedetomidine and buprenorphine but without administering ketamine before the surgery. Our main stimulation paradigm (2 mA pulses administered at 8 Hz) induced seizures in all three SD rats (Figure 4), confirming that ketamine is not necessary for increasing the susceptibility to seizures.

We then used a combination of dexmedetomidine and isoflurane without buprenorphine, and we observed seizures response to our main stimulation paradigm in 2 out of 4 rats (Figure 5). This makes it possible to exclude buprenorphine as an agent necessary for increasing the observed susceptibility to seizures.

Isoflurane has anticonvulsant properties: it suppresses drug-induced convulsions (Kofke et al., 1987) and terminates status epilepticus in patients when administered at concentrations of 0.5%-3.0% (Kofke et al., 1985; Ropper et al., 1986). This makes it unlikely that isoflurane increases the susceptibility to seizures. In addition, we obtained seizures in rats under dexmedetomidine and buprenorphine sedation, without administrating isoflurane, showing that isoflurane is not necessary for the seizure generation.

In Bortel et al. (2019), we applied a similar stimulation paradigm – although we stimulated a single digit, not the entire forepaw. Under dexmedetomidine and buprenorphine sedation, we observed seizures in response to digit stimulation in 6/10 rats. Under urethane anesthesia, with no administration of buprenorphine, we did not observe any seizures.

The only other compounds which we administered in all experiments – those with seizures and those without seizures – are xylazine, which we injected just before the intubation, and carprofen, which we injected approximately at the same time, before the surgical procedures. Similar to dexmedetomidine, xylazine belongs to the α2-adrenoceptor agonists that exert sedation and have additional muscle-relaxant and analgesic properties. A single standard dose of xylazine produces sedation of 30-40 min duration, and its elimination half-life is 30 min (Pawson, 2008). With such elimination half-life, it is very unlikely that xylazine is the compound that increases the susceptibility to seizures in our experiments, because we observed seizures during the time frame of 1 to 3 hours following the initiation of the dexmedetomidine, which is equivalent to 1 ½ to 3 ½ hours after injecting xylazine. Carprofen is a non-steroidal anti-inflammatory drug (NSAID) that has non-narcotic analgesic and antipyretic activity. The mechanism of action of carprofen, like that of other NSAIDs, involves inhibition of cyclooxygenase (COX-2) activity (Rubio et al., 1980). COX-2 is known to play an important role in the early inflammatory response to an insult and in post-seizure inflammation and hyper-excitability of the brain. It was shown that pretreatment of animals with a selective COX-2 inhibitor before applying a convulsant stimulus weakens seizure intensity (Rojas et al., 2014). Therefore, it is unlikely that carprofen increases the susceptibility to seizures in our experiments; in fact, we expect that it decreases this susceptibility. Lastly, we did administer xylazine and carprofen before the surgeries of the experiments we ran with urethane, in which we did not observe any seizures (Bortel et al., 2019). This means that neither xylazine nor carprofen can be the agent causing the seizures.

The results summarized above leave dexmedetomidine as the sole agent that could increase the susceptibility to seizures.

### Adding isoflurane to dexmedetomidine does not prevent the increased susceptibility to seizures

It is widely accepted that anesthetics modify the balance between excitation and inhibition towards increased relative inhibition, and consequently, they modulate evoked neuronal responses (Franceschini et al., 2010). Isoflurane is a general anesthetic that affects many neurotransmitter systems; it acts on GABA, NMDA, and glycine receptors (Grasshoff and Antkowiak, 2006). Although isoflurane reduces neuronal excitation and cerebral metabolism, it is commonly used in electrophysiology studies. However, at higher doses (≥1.6%) isoflurane increases the baseline cerebral blood flow (CBF) (Eger, 1984; Franceschini et al., 2010), and for this reason, it is no longer the anesthetics of choice for hemodynamics-based functional studies.

However, fMRI studies often use a combination of dexmedetomidine and low-percentage of isoflurane in rats (Fukuda et al., 2013; Paasonen et al., 2018) and mouse models (Bukhari et al., 2017; Grandjean et al., 2014). Combining dexmedetomidine with isoflurane constitutes an attractive anesthesia regime for fMRI studies because both cortical and striatal functional connectivity can be reliably detected with no adverse side effects. Moreover, with dexmedetomidine alone, the sedation effect lasts 60-90 minutes even if the administration is continuous, thus forcing short imaging sessions (Pawela et al., 2009). When adding low-percentage isoflurane to dexmedetomidine, the anesthesia effects of isoflurane make it possible to run longer experiments. Importantly, dexmedetomidine and isoflurane provide a synergistic effect as they exert opposing effects on the cerebrovascular system. Whereas isoflurane acts as a vasodilator (Eger, 1984; Franceschini et al., 2010), dexmedetomidine induces cerebral vasoconstriction (Jonckers et al., 2015).

As described in the previous section, isoflurane has anticonvulsant properties (Kofke et al., 1985; Kofke et al., 1987; Ropper et al., 1986). Therefore, isoflurane could be expected to suppress seizure generation when combined with dexmedetomidine. Under dexmedetomidine combined with 0.3%-0.5% isoflurane anesthesia, Fukuda et al. (2013) obtained seizure-free responses to 10 s-long blocks of 1 ms long pulses of 1.5 mA administered at 8 Hz. In contrast, we observed seizures in response to our main stimulation paradigm (2 mA pulses administered at 8 Hz) in 2 out of 4 rats anesthetized with dexmedetomidine combined with 0.3%-0.5% isoflurane. We, therefore, conclude that adding low-percentage isoflurane to dexmedetomidine does not prevent the increased susceptibility to seizures caused by dexmedetomidine. In the next section, we propose stimulation parameters for obtaining seizure-free responses under 3 anesthesia regimes, including the combination of dexmedetomidine and low-percentage isoflurane.

### Blood-oxygenation and cerebral blood flow responses to seizures

The blood-oxygenation and cerebral blood flow responses to seizures induced by forepaw stimulation have a higher amplitude and a larger spatial extent relative to physiological responses to the same stimuli (Figures 2 and 3). It is generally accepted that epileptic events consist of synchronous, rhythmic firing of a population of pathologically interconnected neurons capable of generating high-frequency oscillations (Bragin et al., 2002; Shariff et al., 2006). Each physiological increase in neuronal activity increases the cerebral metabolic rate of oxygen consumption, leading to an increase in CBF and CBV as the brain attempts to perfuse sufficiently the active neurons with oxygenated hemoglobin (Schwartz and Bonhoeffer, 2001; Shariff et al., 2006; Zhao et al., 2008). The signaling molecules such as adenosine and nitric oxide are released by the firing neurons, causing nearby pial arterioles to dilate (Haglund and Hochman, 2007). Consequently, seizures elicit a large focal increase in metabolism and utilization of oxygen and glucose, resulting in an enormous increase in blood volume and flow to the ictal focus to provide adequate oxygenation (Engel et al., 1982; Harris et al., 2018; Patel et al., 2013). It was demonstrated that during pilocarpine-induced status epilepticus a compensation phase lasting up to 30 min is observed with an acute increase in CBF, followed by a decompensation phase with CBF decrease (Choy et al., 2010; Lothman, 1990; Reddy and Kuruba, 2013). There is a preferential distribution of blood flow to certain regions of the brain during seizures. The degree of perfusion change in the cortex is greater than in the thalamus, and the hippocampus is hypo-perfused when compared to the cortex (Choy et al., 2010). In the present experiment, an increase in cerebral hemodynamics is observed during seizures simultaneously with an increase in neuronal activity, extending beyond the period and spatial extent of normal responses to sensory stimulation.

The findings reported here are in agreement with previous studies that characterized the cerebral hemodynamics during seizures: partial seizures have widespread effects on cortical function and cerebral perfusion (Harris et al., 2014; Zhao et al., 2007).

### Seizure-free stimulation paradigms and anesthesia regimes for functional imaging in rats

Dexmedetomidine alone and dexmedetomidine combined with low-percentage isoflurane are two commonly used anesthesia regimes for functional imaging. However, our results show that these anesthesia regimes increase the susceptibility to seizures in rats. We observed seizures when applying forepaw stimulation of 10 s blocks with 1 ms long pulses of 1.0-2.0 mA administered at 8 Hz (Figures 1–5).

Similar potent peripheral stimulation paradigms in the range of 8-12 Hz and 1-2 mA are commonly used to elicit BOLD responses in dexmedetomidine sedated rats (Huttunen et al., 2011; Kim et al., 2010; Nunes et al., 2019; Paasonen et al., 2017; Schulz et al., 2012). These studies likely induced seizures and interpreted fMRI responses to seizures as normal responses.

We, therefore, conducted experiments to determine the stimulation parameters for seizure-free experiments under these anesthesia regimes. Our results show that when using relatively long stimulations blocks of 10 s – that are required for fMRI block paradigms – forepaw stimulation with 1 ms long pulses of 1.5–2.0 mA at 4 Hz elicits seizure-free responses (Figures 6 and 9). Importantly, these stimulation parameters elicit hemodynamic responses that can be clearly detected in single trials (Figures 9 and 10). Therefore, we propose these paradigms for safe stimulation that does not induce seizures but still generates well detectable hemodynamic responses.

Another possibility is to use urethane anesthesia, in studies that do not require recovery of the animal following the neurophysiology and/or functional imaging experiment. Urethane has modest effects on both the inhibitory and excitatory systems and does not affect the noradrenergic system (Hara and Harris, 2002). The changes it exerts on multiple neurotransmitter-gated ion channels are much smaller than those seen with anesthetics more selective for one neurotransmitter system, such as ketamine. Therefore, urethane is suitable for maintaining anesthesia during electrophysiological recording and functional imaging (Hara and Harris, 2002). As we demonstrated previously (Bortel et al., 2019), under urethane anesthesia, 10 s blocks of digit stimulation with 1 ms long pulses of 2 mA administered at 8 Hz elicited seizure-free, well detectable neurophysiological and hemodynamic responses.

### Mechanisms of stimulation-induced seizures under dexmedetomidine sedation

Both sedation and general anesthesia suppress the central nervous system, but only general anesthesia results in unconsciousness and lack of sensation (Miller, 2010; Turner and Knapp, 1995; Young-McCaughan and Miaskowski, 2001). Under general anesthesia, the activity of the thalamic and midbrain reticular formation nuclei is suppressed. The suppression of the thalamic information transfer disrupts the somatosensory input from reaching higher cortical areas (Miller, 2010). Dexmedetomidine has sedative properties, and buprenorphine is an opioid with analgesic properties. It is possible that the combination of dexmedetomidine and buprenorphine modifies the anesthesia regime from sedation (with dexmedetomidine alone) closer to anesthesia (with dexmedetomidine and buprenorphine). However, since this is unclear, here we use the term ‘sedation’ when describing the effect of dexmedetomidine and buprenorphine. In contrast, given that isoflurane is a general anesthetic, we use the term ‘anesthesia’ when referring to the effect of combining dexmedetomidine and isoflurane.

Dexmedetomidine is an α2-adrenergic agonist with sedative properties, predominantly acting on presynaptic receptors in the locus coeruleus. It regulates the central adrenergic function and in consequence induces cerebral vasoconstriction mediated by direct agonist binding to receptors on the cerebral vessels, resulting in a reduced baseline of CBF and CBV (Adamczak et al., 2010; Jonckers et al., 2015; Paasonen et al., 2018; Pawela et al., 2009; Weber et al., 2006). The degree of vasoconstriction depends on the dose and delivery method (topical vs. systemic) (Jonckers et al., 2015). Dexmedetomidine decreases noradrenaline release (Gertler et al., 2001), which in turn decreases seizure threshold (Oishi and Suenaga, 1982).

There have been several conflicting reports on the influence of dexmedetomidine on seizure generation. It was shown that the seizure frequency and onset time evoked by kainic acid (Airaksinen et al., 2012; Airaksinen et al., 2010) or pilocarpine (Choy et al., 2010) are similar in the awake and medetomidine-sedated rats. Halonen et al. (1995) demonstrated that rat convulsions induced by kainic acid are prevented by the administration of medetomidine. Similarly, it was shown that rat seizures induced by the administration of cocaine were suppressed by medetomidine sedation (Whittington et al., 2002). Furthermore, high doses (100 μg/kg or higher) of dexmedetomidine effectively decreases the number and cumulative time of repeated seizures evoked by prolonged intracranial electrical stimulation of the amygdala (Kan et al., 2013). In contrast, it was shown that dexmedetomidine increases the epileptiform activity in epileptic patients (Chaitanya et al., 2015). In addition, dexmedetomidine exerts a significant proconvulsant action in the pentylenetetrazol seizure animal model. The proconvulsant effect is dose-dependent and stereospecific. It can be blocked by the selective α2-adrenergic antagonist atipamezole (Mirski et al., 1994). Likewise, Fukuda et al. (2013) demonstrated that epileptic activity could be induced in rats by electrical stimulation of the forelimb after 2 hours-long continuous IV infusion of dexmedetomidine (Fukuda et al., 2013).

In line with the findings by Whittington et al. (2002) and Airaksinen et al. (2012), we did not observe any seizure-like responses to short and weak forelimb stimuli (1 s-long, 0.6-0.8 mA current pulses delivered at 8 Hz) under dexmedetomidine sedation (Sotero et al., 2015). Here, we did not observe seizures with 10 s-long stimulation with either 1 mA or 2 mA pulses delivered at 4 Hz (Figure 6A-B), as was also shown by Fukuda et al. (2013). Nevertheless, under the same sedation regime, long (10 s), potent (1 mA or 2 mA), repetitive and frequent (8 Hz) peripheral electrical stimuli evoke seizures (Figures 1-5 and 7A-C), as was also demonstrated by Bortel et al. (2019). Therefore, dexmedetomidine increases the susceptibility to seizures. However, potent stimuli are necessary, too, for inducing seizures by peripheral stimulation under such increased susceptibility.

We did not perform systematic experiments to test the effect of dexmedetomidine dose on the susceptibility to seizure. Therefore, we cannot rule out the possibility that the susceptibility to the generation of seizures depends on the dose of dexmedetomidine. However, we obtained data that indicate that the lack of seizures in C57 mice did not depend on the dose. To maintain proper sedation in three of the mouse experiments, we had to increase the rate of dexmedetomidine from 0.05 mg/kg/h to 0.1 mg/kg/h approximately 80 minutes after the infusion started. However, no seizures were observed in these mice or the other mice. These results are consistent with Fukuda et al. (2013)’s findings that increasing the dexmedetomidine IV infusion rate from 0.05 mg/kg/h to 0.15 mg/kg/h in SD rats does not change the susceptibility to seizure generation.

The trigger to the epileptic activity in our animals is repetitive electrical somatosensory stimulation. Therefore, these seizures are reflex seizures. Reflex seizures are defined as seizures triggered by repetitive, 5-20 s long sensory stimulations (Kanemoto et al., 2001; Panayiotopoulos, 2005; Sala-Padro et al., 2015; Striano et al., 2012; Wolf, 2015). Epileptic activity induced in our rats consisted of brief and focal electrographic seizures; the animals did not show any epileptic behavior. Similar to our observation in rats, touch induced seizures in humans manifest as focal, brief seizures that may have only an electrographic display without any overt clinical manifestations (Panayiotopoulos, 2005).

### Dexmedetomidine sedation increases susceptibility to seizures in rats but not in wild-type mice

We have shown that dexmedetomidine increases susceptibility to seizures in SD and LE rats, but not in C57 mice. Previous works have demonstrated that the rat and mouse strains show diverse susceptibility to seizure-induction (Ferraro et al., 1995; Golden et al., 1995; McKhann et al., 2003). Wistar-Furth rats are more sensitive to the convulsant effects of kaininc acid (KA) than Sprague-Dawley and Long-Evans Hooded rats (Golden et al., 1995; Golden et al., 1991). It was shown that the C57BL6 mouse strain has lower seizure sensitivity than several strains reported as sensitive to seizures, including the SWR/J, FVB/NJ, CBA/J, DBA/1, and DBA/2 strains (Frankel et al., 2001). Indeed, DBA/2J mice exhibit a higher susceptibility to maximal electroshock and KA-induced seizures relative to C57BL/6J mice (Ferraro et al., 1995; Ferraro et al., 2002). Similarly, McKhann et al. (2003) showed that C57BL/6J and C3HeB/FeJ mice are more resistant to seizures than 129/SvEms mice. Our results support the assertion that C57 mice have low susceptibility to seizures. More in general, our findings support the concept that seizure susceptibility depends on the animal’s species and strain.

Our findings demonstrate the lack of significant, sustained LFP responses to 8 Hz — and only reduced LFP responses to 6 Hz — relative to responses to 4Hz forepaw stimulation in mice (Fig. 12A-C). The maintenance of the proper balance between excitation and inhibition is critical for the normal function of cortical circuits (Turrigiano, 2011). This continuum of changes in response amplitude suggests that changes in the balance between excitation and inhibition take place as a function of the stimulation frequency. We propose that the inhibitory tone is overall enhanced at higher stimulation frequencies, possibly because the inhibitory response following a stimulus is still in effect when the next stimulus is administered in a train of stimulation at 8 Hz but not at 4 Hz. Indeed, one of the explanations proposed by Wang et al. (2013) is that the effect of inhibition remains longer than that of the excitation, and therefore the subsequent excitation in a train overlaps the inhibition. It was also shown that a decrease in the firing of neurons to high-frequency stimuli is a consequence of an altered balance in excitatory and inhibitory cortical activity (House et al., 2011; Li et al., 2009).

Our findings in mice experiments differ from Fukuda et al. (2013)’s findings and our findings (Figures 6 and 7) in SD rat experiments. Fukuda et al. (2013) reported that under dexmedetomidine sedation (with the addition of pancuronium bromide) the evoked LFP responses to forepaw stimulation were larger at a frequency of 8–10 Hz than those elicited by 4 Hz, 6 Hz, and 12 Hz stimulation. The difference in the stimulus frequency dependence of the LFP responses between mice and SD rats suggests that the effect of stimulus frequency on the balance between excitation and inhibition is different for these two species.

## CONCLUSION

Our findings demonstrate that hemodynamic responses to potent stimuli in rats sedated with dexmedetomidine or anesthetized with dexmedetomidine and isoflurane are possibly due to induced seizures. We ruled out the possibility that the increased susceptibility to seizures in our experiments was caused by any agent other than dexmedetomidine. The induction of seizures in experiments that use dexmedetomidine depends on stimulation strength. To obtain physiological yet detectable neuronal and hemodynamic responses, we propose stimulation parameters that ensure seizure-free responses under dexmedetomidine sedation or dexmedetomidine and isoflurane anesthesia. By definition, obtaining spontaneous activity for the study of the resting-state condition does not involve administering stimuli. Indeed, we did not observe any seizures during spontaneous activity (data not shown).

Our results reveal that dexmedetomidine increases susceptibility to seizures in the 2 rat strains we tested, SD and LE rats, but not in wild type C57 mice. More in general, our findings support the concept that seizure susceptibility depends on the animal’s species.

## Acknowledgments

This study was supported by the Canadian Institute of Health Research (Grant MOP-102599), and the Natural Sciences and Engineering Research Council of Canada (RGPIN 375457-09 and RGPIN 2015-05103) awarded to A.S. We thank Jean Gotman, Massimo Avoli, Victor Mocanu and Pascal Kropf for their helpful technical advice and discussions.

